# Development of recombinant monoclonal antibodies targeting conserved VlsE epitopes in Lyme disease pathogens

**DOI:** 10.1101/2022.05.05.490853

**Authors:** Li Li, Lia Di, Saymon Akther, Brian M. Zeglis, Weigang Qiu

**Affiliations:** Graduate Center, City University of New York, NY, USA; Department of Biological Sciences, Hunter College, City University of New York, NY, USA; Department of Chemistry, Hunter College, City University of New York, NY, USA; Department of Radiology, Weill Cornell Medical College, NY, USA; Department of Radiology, Memorial Sloan Kettering Cancer Center, NY, USA; Department of Physiology and Biophysics & Institute for Computational Biomedicine, Weill Cornell Medical College, NY, USA

## Abstract

VlsE (<u>v</u>ariable major protein-<u>l</u>ike <u>s</u>equence, <u>e</u>xpressed) is an outer surface protein of the Lyme disease pathogen (*Borreliella* species) and a key diagnostic biomarker of Lyme disease. However, the high sequence variability of VlsE poses a challenge to the development of consistent VlsE-based diagnostics and therapeutics. In addition, the standard diagnostic protocols detect immunoglobins elicited by the Lyme pathogen, not the presence of pathogen or its derived antigens. Here we describe the development of recombinant monoclonal antibodies (rMAbs) that bind specifically to conserved epitopes on VlsE. We first quantified amino-acid sequence variability encoded by the *vls* genes from thirteen *B. burgdorferi* genomes by evolutionary analyses. We showed broad inconsistencies of the sequence phylogeny with the genome phylogeny, indicating rapid gene duplications, losses, and recombination at the *vls* locus. To identify conserved epitopes, we synthesized peptides representing five long conserved invariant regions (IRs) on VlsE. We tested the antigenicity of these five IR peptides using sera from three mammalian host species including human patients, the natural reservoir white-footed mouse (*Peromyscus leucopus*), and VlsE-immunized New Zealand rabbits (*Oryctolagus cuniculus*). The IR4 and IR6 peptides emerged as the most antigenic and reacted strongly with both the human and rabbit sera, while all IR peptides reacted poorly with sera from natural hosts. Four rMAbs binding specifically to the IR4 and IR6 peptides were identified, cloned, and purified. Given their specific recognition of the conserved epitopes on VlsE, these IR-specific rMAbs are promising diagnostic and theragnostic agents for direct detection of Lyme disease pathogens regardless of strain heterogeneity.

## Introduction

Lyme disease is a multistage, tick-transmitted infection caused by spirochetes of the bacterial species complex *Borrelia burgdorferi sensu lato* (*Bbsl*), known more concisely (albeit controversially) as a new genus *Borreliella* (Barbour and Qiu, 2019; Margos *et al*., 2020). Lyme disease is the most common tick-borne disease in regions of North America, Europe, and Asia (Stanek *et al*., 2012; Kilpatrick *et al*., 2017). In the United States, approximately 476,000 cases are diagnosed annually (Kugeler *et al*., 2021). The majority of Lyme disease cases in the US are caused by the single species *B. burgdorferi* and transmitted by the hard-bodied *Ixodes scapularis* or *I. pacificus* ticks, although the same tick vectors carry other *Borreliella* species as well as *Borrelia* species closely related to relapsing fever spirochetes (Barbour *et al*., 2009; Pritt *et al*., 2016; Schwartz *et al*., 2021). *B. burgdorferi* cause multisystemic manifestations in humans including erythema migrans (EM) at early stages, arthritis, carditis, and neuroborreliosis in late stages, and chronic symptoms associated with persistent infections (Stanek *et al*., 2012; Sharma *et al*., 2015; Feng *et al*., 2019).

Antigenic variation via continuously altering the sequences of surface antigens during infection is a common strategy that microbial pathogens employ to escape adaptive immune responses of vertebrate hosts (Vink, Rudenko and Seifert, 2012; Palmer, Bankhead and Seifert, 2016). In the two closely spirochetal groups of *Borrelia* causing relapsing fever and *Borreliella* causing Lyme disease, two homologous but distinct molecular systems have evolved facilitating continuous antigenic variation through recombination between an expressed locus and silent archival loci during persistent infection within the vertebrate hosts (Norris, 2006). In *B. burgdorferi*, the molecular system able to generate antigenic variation consists of one expression site (*vlsE*, variable major protein-like sequence, expressed) and a set of tandemly arranged silent cassettes (*vlsS*) that share more than 90% similarities to the central cassette region of *vlsE* (Zhang *et al*., 1997; Norris, 2014; Verhey, Castellanos and Chaconas, 2019) (Fig 1). During mammalian infection, *vlsE* continuously expresses and undergoes random segmental recombination with the silent cassettes, generating a considerable number of new VlsE antigen variants to prolong spirochete infection in hosts (Norris, 2006; Verhey, Castellanos and Chaconas, 2019).

**Fig 1.**
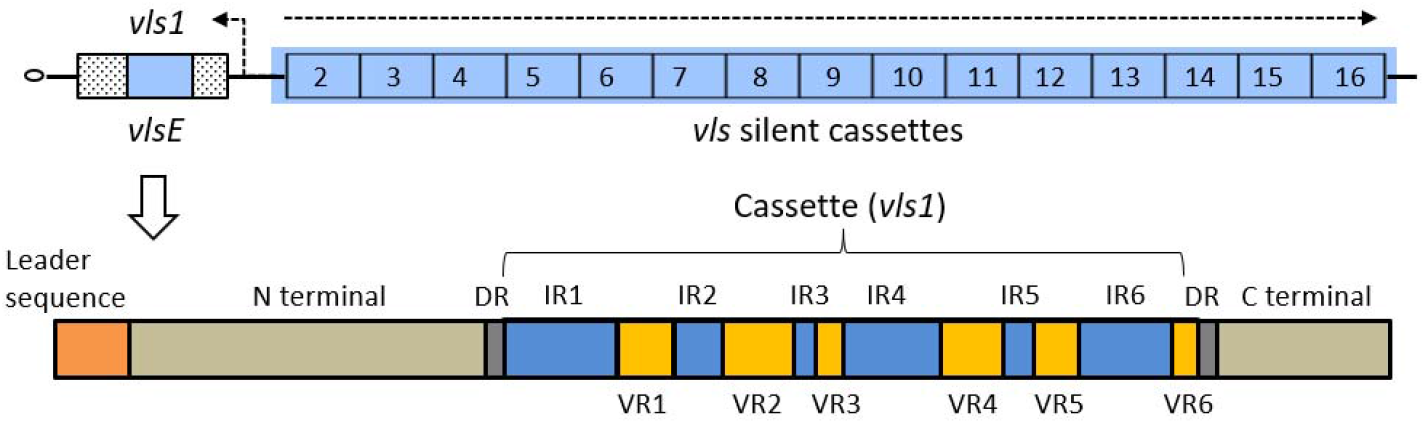
Genomic and gene structures of the *vls* locus in *B. burgdorferi* strain B31 (A) The *vls* locus is located close to the telomere of the linear plasmid lp28-1 (GenBank accession AE000794) in the B31 genome, consisting of cassettes of silent (un-expressed) open reading frames (ORFs) (*vls2* through *vls16*) and an expressed ORF (*vlsE*) containing the *vls1* cassette introduced by recombination (Zhang *et al*., 1997). Dashed arrows indicate the direction of coding strands. (B) The VlsE protein consists of a leader peptide, a N-terminus domain, a cassette flanked by two direct repeats (DRs), and a C-terminus domain. The central cassette consists of interspersed variable (VR1-6) and invariant regions (IR1-6).

The *vlsE* gene encodes a 36 kD lipoprotein that is anchored to the outer membrane on the cell surface. The primary structure of VlsE comprises the N- and C-terminal domains as well as the central cassette which consists of six highly variable regions (VR1-VR6) interspersed with six conserved invariant regions (IR1-IR6) (Fig 1). The N- and C-terminal regions do not undergo antigenic variation and are thought to be important in maintaining the functional structure of the molecule (Norris, 2014). The VRs of the cassette are the sequences that undergo antigenic variation during infection, while the IRs are conserved among *B. burgdorferi* strains (Liang *et al*., 1999). The crystal structure of recombinant VlsE protein revealed that the six VRs constitute loop structures and form a “dome” on the membrane distal surface exposed to the host environment, which may shield the IRs from antibody binding (Eicken *et al*., 2002).

VlsE elicits strong humoral responses that can be detected throughout the course of Lyme disease, making it a powerful antigen in serologic assays of Lyme disease diagnosis (Lawrenz *et al*., 1999; Bacon *et al*., 2003; Elzbieta *et al*., 2016). Contrary to the established paradigm of weak immunogenicity of the conserved regions of bacterial surface proteins, the conserved IR6 elicits immunodominant antibody responses during human infection despite the region being largely inaccessible on the intact VlsE molecule (Liang *et al*., 1999; Chandra *et al*., 2011; Elzbieta *et al*., 2016). The surprising finding of immunodominance of IR6 in human patients is hypothesized to be a result of antigen processing of the VlsE proteins in non-reservoir host species (Embers *et al*., 2007).

A 26-amino acid peptide that reproduces the IR6 sequence, known as C6 peptide, is used in commercial diagnostic tests of Lyme disease (Bacon *et al*., 2003; Wormser *et al*., 2013). The standardized two-tiered testing (STTT) for Lyme disease diagnosis includes a screening enzyme immunoassay (EIA) with the whole cell sonicate and a subsequent confirmatory Western blot assay for the presence of both IgM and IgG antibodies against ten *Borreliella* antigens (CDC, 1995; Moore *et al*., 2016). Recently, a modified two-tiered testing (MTTT) protocol using two sequential EIAs with the C6 peptide or the whole VlsE protein has been developed. MTTT improved sensitivity and specificity relative to STTT, especially in Lyme patients with early-stage manifestations (Branda *et al*., 2017). Nevertheless, the overall sensitivity for early-stage diagnosis remains low, ranging from 36% to 54%, even with MTTT (Pegalajar-Jurado *et al*., 2018). In addition, both diagnostic assays are indirect tests and do not distinguish between active infection and past exposure. In sum, there is a need to simplify the testing protocol for Lyme disease, improve testing sensitivity in the early infection stage, and detect the presence of Lyme pathogen or its derivative antigens directly.

During the transmission cycle of *B*. burgdorferi, the *vls* locus is expressed during the late-stage persistent infection within the mammalian host, in contrast to genes like *ospA* (encoding outer surface protein A) expressed within the ticks, and genes like *ospC* expressed exclusively during a short window of time when the spirochetes begin migrate from the tick to the mammalian host (Samuels, 2011; Tilly, Bestor and Rosa, 2013). As a multi-copy gene family and driven by adaptive amino-acid substitutions, the *vls* cassettes exhibit high sequence variability not only between *B. burgdorferi* strains but also within the same genome (Glöckner *et al*., 2004; Schutzer *et al*., 2011; Graves *et al*., 2013). In the present study, we developed a bioinformatics workflow to facilitate the automated identification of *vls* sequences from the sequenced *Borreliella* genomes. We quantified evolutionary rates at individual amino acid sites of the *vls* coding sequences identified from thirteen *B. burgdorferi* genomes. Extending the previous analysis of mechanisms of evolution at the *vls* locus (Graves *et al*., 2013; Schwartz *et al*., 2021), we explored the evolution mechanisms by comparing the *vls* gene phylogeny with the genome-derived strain phylogeny. Our experimental investigations of the immunogenicity of the VlsE protein confirmed the immunodominance of the IR6 peptide and discovered the similar immunodominance of the IR4 peptide in human patients and immunized rabbits but not the reservoir hosts. Finally, we identified, cloned, and purified four recombinant IR-specific monoclonal antibodies (rMAbs) that are promising theragnostic agents for direct assay of *B. burgdorferi* infection in clinical samples and model organisms of Lyme disease.

## Materials and Methods

### Identification of *vls* cassette sequences and evolutionary analysis

We downloaded the whole genome sequences of 12 *B. burgdorferi* strains from NCBI GenBank (Schutzer *et al*., 2011). The *vlsS* sequences of B31-5A3 clone (GenBank accession U76406) (Zhang *et al*., 1997) were used as the queries to searched for sequences homologous to the *vls* cassette sequences using HMMER (version 3.3.2) (Eddy, 2011). A customized web-based software tool was developed to identify and extract individual *vls* sequences given a *B. burgdorferi* replicon sequence (http://borreliabase.org/vls-finder). The identified *vlsS* and *vlsE* sequences were translated, aligned, and converted into a codon alignment using MUSCLE (version 3.8.31) (Edgar, 2004) and the *bioaln* utility (--*dna2pep* method) of the BpWrapper (version 1.13) toolkit (Hernández *et al*., 2018). A maximum likelihood tree was subsequently inferred using IQ-TREE (version 1.6.1) with the best-fit nucleotide substitute model KOSI07 and 1000 bootstrap replicates (Nguyen *et al*., 2015). Branches with lower than 80% bootstrap support were collapsed using the *biotree* utility (*-D* method) of the BpWrapper utility (Hernández *et al*., 2018). The tree was rendered using the R package *ggtree* (Version 2.2.4) (Yu *et al*., 2017). To quantify the sequence conservation, evolutionary rates at individual amino acid positions were estimated using Rate4Site (version 3.0.0) with the protein alignment and the phylogenetic tree as inputs and the B31-5A3 VlsE sequence (GenBank accession U76405) as the reference (Pupko *et al*., 2002). Sequence conservation at the IRs was further quantified and visualized with WebLogo (Crooks *et al*., 2004).

### Synthesis of peptides representing conserved epitopes of VlsE

The preparation of the peptides was based on the annotation of B31-5A3 VlsE protein sequence in the literature (Zhang *et al*., 1997). Five invariant regions, IR1, IR2, IR4, IR5, and IR6, were tested for antigenicity using sera from three host species. IR3 (AGKLFVK), the shortest IR, was excluded from the antigenicity test. Extra flanking amino acids were added to IR2, IR4, and IR5 to meet the minimum length for peptide synthesis. Peptides were commercially synthesized and biotin-labeled on the N-terminus using Fmoc chemistry (GenScript, Piscataway, NJ, USA). Sequences of these peptides are shown in Table 1.

**Table 1.**
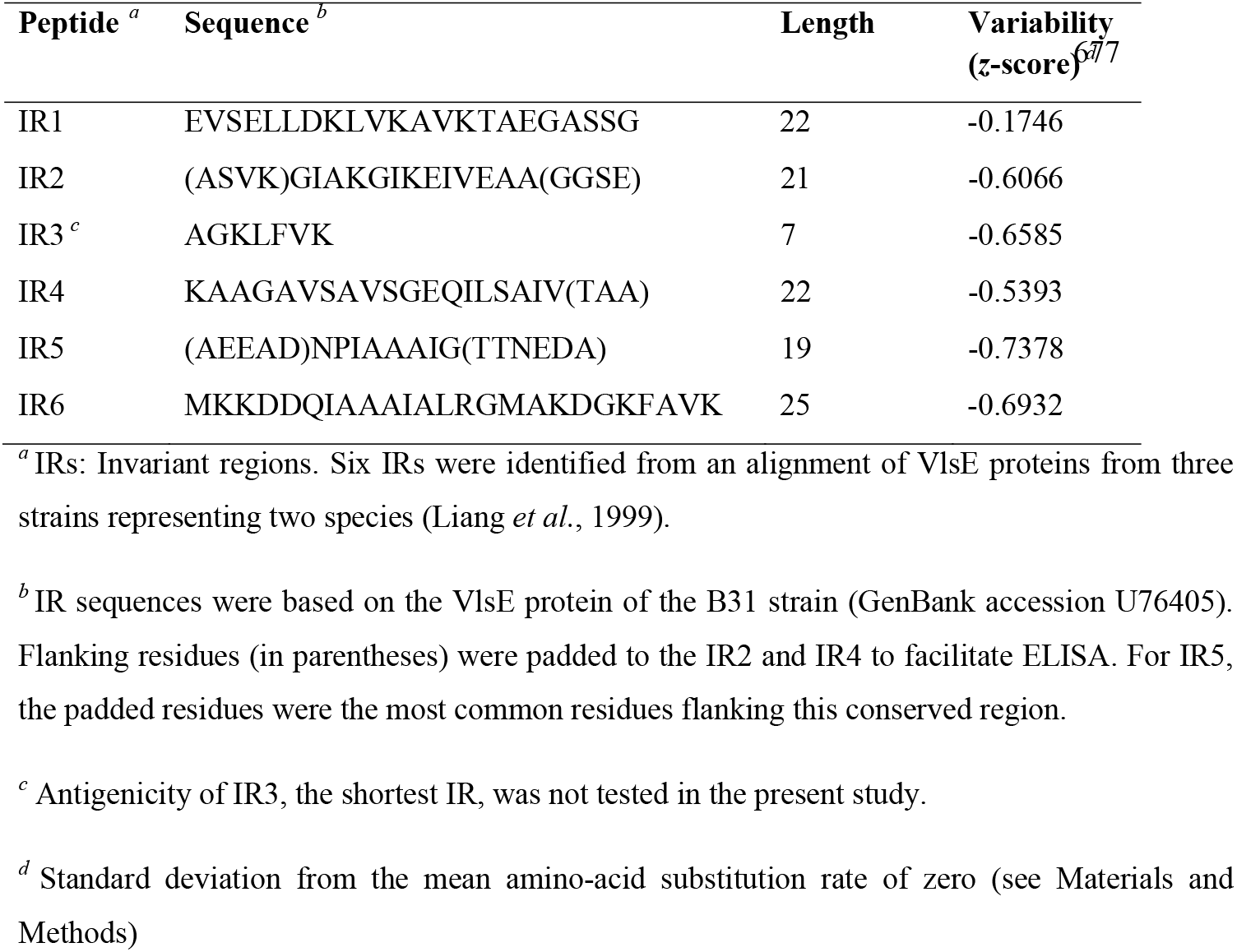
Peptides used to screen for IR-specific monoclonal antibodies

**Table 2.**
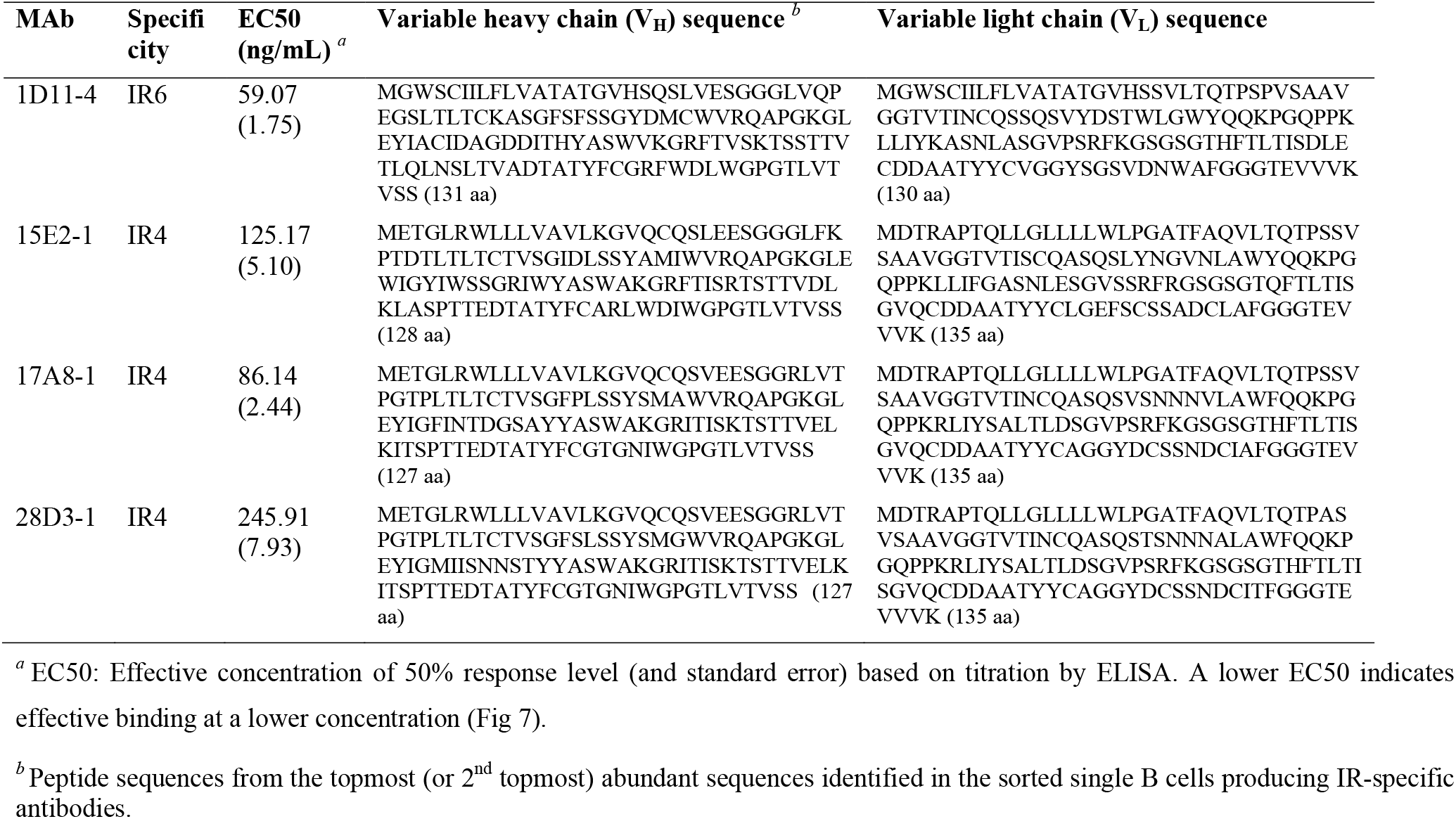
Specificity and sequences of IR-specific mAbs

### Sera collection from naturally infected hosts

The 56 serum samples, consisting of Lyme patient and control sera provided by the US Center for Disease Control and Prevention (CDC, *n* = 40), Lyme patient sera provided by Dr Maria Gomes-Solecki (University of Tennessee Health Science Center, *n* = 6), and sera from the reservoir hosts (white-footed mouse, *Peromyscus leucopus*) provided also by Dr Maria Gomes-Solecki (*n* = 10), have been used and described in previous publications (Ivanova *et al*., 2009; Molins *et al*., 2014; Di *et al*., 2021). Briefly, among the human samples, 25 serum samples were derived from patients with early-stage Lyme disease including those diagnosed as having the skin symptom erythema migran (EM) or as EM convalescence. Seventeen human sera samples were from patients displaying late-stage Lyme disease symptoms including arthritic, cardiac, and neurological Lyme diseases. Four human sera samples were from healthy individuals as controls.

### Cloning, over-expression, and purification of recombinant VlsE protein

Recombinant VlsE protein from the B31 strain was cloned, over-expressed, and purified using a protocol described preciously (Di *et al*., 2021). Briefly, the 585-bp *vlsE* cassette region (including the direct repeat regions on both ends) of B31-5A3 clone was codon-optimized, synthesized, and cloned into the pET24 plasmid vector which then transfected *Escherichia coli* BL21 cells. A10 × Histidine-tag was added on the N-terminus of the construct to facilitate the downstream purification. All cloning work was performed by a commercial service (GeneImmune Biotechnology Corp., Rockville, MD, USA). The *E. coli* strain that contained a cloned *vlsE* cassette was cultured in Luria-Bertani (LB) broth containing 0.4% glucose and 50 μg/ml Ampicillin. When the culture reached exponential growth, we induced the expression of the cloned *vlsE* cassette by adding isopropyl β-d-1-thiogalactopyranoside (IPTG) to a final concentration of 0.25 mM and by incubation overnight at 25 °C. Cells were collected and then lysed by lysozyme and sonication. The lysate supernatant, containing the recombinant VlsE protein, was purified using nickel sepharose beads (Ni-NTA, Thermo Fisher Scientific, Waltham, MA, USA) following the manufacture’s protocol. The identity and concentration of the purified protein was examined and quantified using the sodium dodecyl sulphate polyacrylamide gel electrophoresis (SDS-PAGE) and the Pierce Bradford Protein Assay Kit (Thermo Fisher Scientific, Waltham, MA, USA).

### Immunization of rabbits and preparation of polyclonal and monoclonal antibodies

Antibody preparation was conducted with a commercial service GenScript (Piscataway, NJ, USA). Briefly, the project consisted of four stages. In Stage 1, animals were immunized and polyclonal antibodies were obtained. Specifically, four New Zealand rabbits (*Oryctolagus cuniculus*) were immunized with 100 μg purified recombinant VlsE protein on Days 1, 14, and 28. The rabbits were bled for antiserum collection one week after the third immunization. The antisera were subsequently purified by affinity chromatography to obtain polyclonal antibodies (pAbs), which were assayed for anti-VlsE activity. In Stage 2, monoclonal antibodies (MAbs) were identified via single B cell sorting. Peripheral blood mononuclear cells (PBMC) were collected from the two selected immunized rabbits one week after a booster dose with the purified VlsE. Plasma B cells (CD138+) were isolated and enriched using a commercial kit. B cells were then screened for VlsE-specific cell lines using ELISA. The supernatants of positive cell lines were used to test for binding with VlsE and positive cell lines were chosen for mAb production. In Stage 3, the variable domains of the light and heavy chains of the VlsE-binding antibodies were sequenced. Total RNA was isolated from the VlsE-binding B cell lines and reverse-transcribed into cDNA using universal primers. Antibody fragments of the heavy chain and the light chain were amplified and sequenced. In Stage 4, the recombinant MAbs (rMAbs) were produced. The amplified antibody variable fragments were cloned into plasmid vector pcDNA3.4 which then transfected mouse cells for expression. Supernatants of cell cultures were harvested continuously. The rMAbs were purified using the Protein A/G affinity chromatography (with immobilized Protein A and G from *Staphylococcus aureus*) followed by size exclusion chromatography (SEC).

### Identification of IR-specific mAbs with ELISA

We tested sera from naturally infected hosts for reactivity to the IR peptides (Table 1) and recombinant VlsE protein with ELISA using a protocol described preciously (Di *et al*., 2021). Briefly, a 96-well MICROLON 600 plate (USA Scientific, Inc., Ocala, FL, USA) was incubated with 10 μg/ml of antigen overnight at 4 °C. Serum samples diluted between 1:100 to 1:1000 were applied after blocking with 5% milk and were incubated for 2 h at 37 °C, followed by application of horseradish peroxidase (HRP)-conjugated secondary antibodies. We used the Goat Anti-Human IgG/IgM (H + L) (Abcam, Cambridge, UK) 1:40,000 for assays of human sera and the Goat Anti-*P. leucopus* IgG (H + L) (SeraCare Life Sciences, MA, USA) 1:1000 for assays of *P. leucopus* sera. The antigen-antibody reaction was probed by TMB ELISA Substrate Solution (Invitrogen eBioscience) and was terminated with 1M sulfuric acid after 15 minutes. Binding intensities were measured at the 450 nm wavelength using a SpectraMax i3 microplate reader (Molecular Devices, LLC, CA, USA).

The purified anti-VlsE pAbs, the supernatants of selected B cell cultures, and the purified mAbs were tested for reactivity to the IR peptides and the purified recombinant VlsE protein with ELISA using the same protocol as described above. Mouse Anti-Rabbit IgG Fr secondary antibody (GenScript, Piscataway, NJ) 1:30,000 was used for assays of these rabbit-derived samples. Serial dilutions of MAbs by factors from 1,000 to 512,000 were tested with ELISA to quantify the binding activities.

### Protein structure visualization

The PDB file of VlsE protein structure (accession 1L8W) was downloaded from the protein data bank (PDB) (https://www.rcsb.org/) (Eicken *et al*., 2002). The PDF file describes a tetramer of VlsE. We used Chimera (version 1.15) (Pettersen *et al*., 2004) to visualize the protein structure in ribbon and surface-filled formats and to color the six invariable regions (IR1-6).

### Animal care, data visualization, statistical analysis, and data and code availability

Antibody production from the New Zealand rabbits followed the protocols approved by the Office of Laboratory Animal Welfare (OLAW) Assurance and the Institutional Animal Care and Use Committee (IACUC) of the vendor (GenScript, Piscataway, NJ).

Data visualization and statistical analysis were performed in the R statistical computing environment (R Core Team, 2013) accessed with RStudio. The alignment of translated *vls* sequences, ELISA readings, and R scripts are publicly available on Github at https://github.com/weigangq/vls-mabs.

## Results

### Phylogenetic inconsistencies indicate duplications and losses, sequence divergence, and recombination at the *vls* locus

We identified 194 *vls* cassette sequences from 13 *B. burgdorferi* strains and inferred a maximum likelihood tree of the cassette (Fig 2). These *B. burgdorferi* strains have been classified into four phylogenetic groups (A-D) based on chromosomal single-nucleotide polymorphisms (SNPs) (Mongodin *et al*., 2013). The *vls* gene phylogeny consists of eight major clades and is consistent with a previously published *vls* cassette phylogeny (Graves *et al*., 2013). Here we analyzed the *vls* gene phylogeny in the broader context of strain phylogeny. Phylogenetic inconsistencies between gene and strain trees may result from – and thus indicate the occurrence of – horizontal gene transfers between strains, ancestral gene duplications followed by the loss of duplicated copies, and incomplete lineage sorting when strains rapidly diverge from one other (Rogers *et al*., 2017; Kundu and Bansal, 2018).

**Fig 2.**
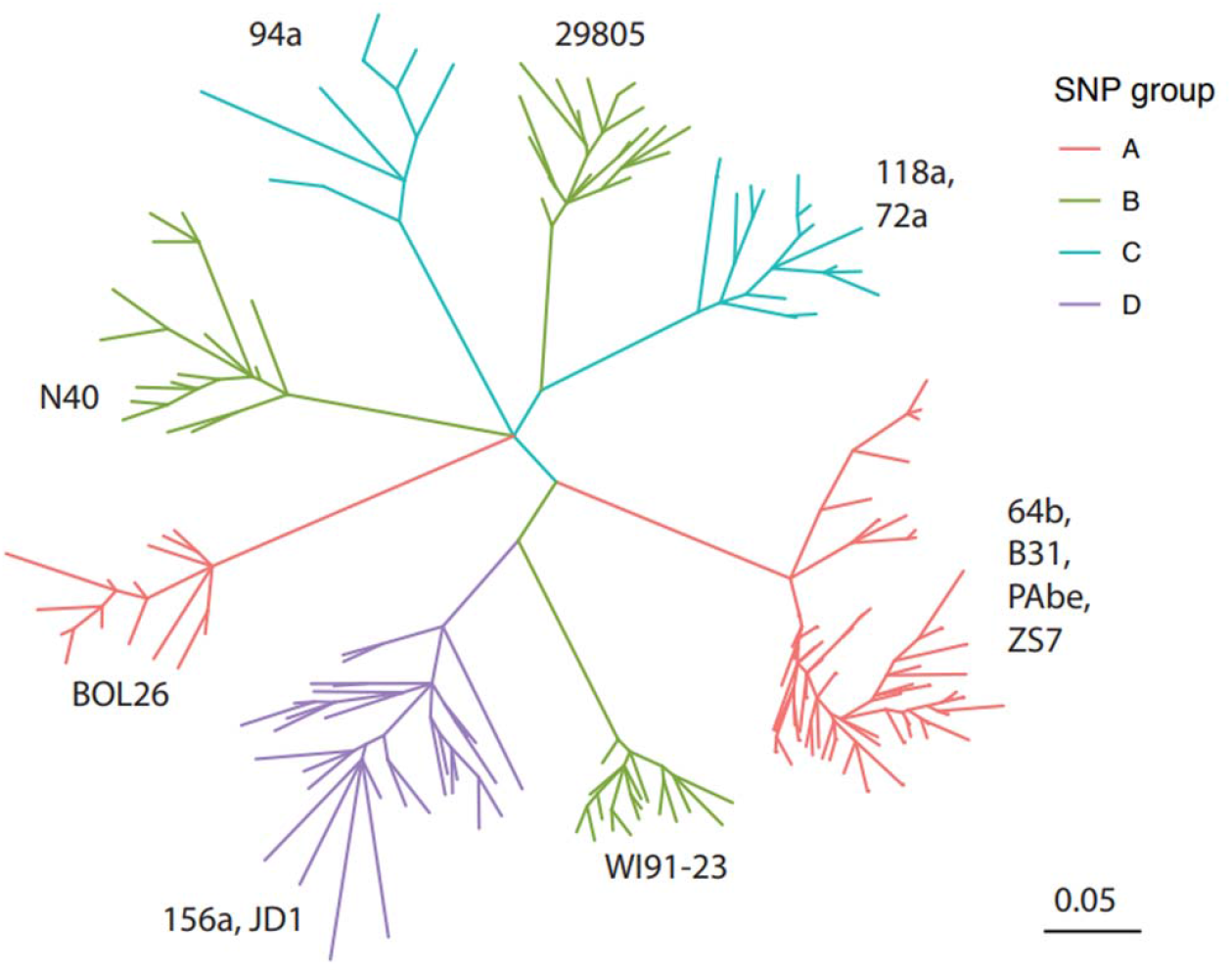
Sequence diversity of *vls* cassettes of 13 *B. burgdorferi* strains Eight major clades of *vls* alleles were identified based on the codon alignment of 194 cassette sequences from 12 US *B. burgdorferi* genomes (strain names shown by the clades) (Schutzer *et al*., 2011). The maximum likelihood tree was inferred using IQ-TREE (version 1.6.1) (Nguyen *et al*., 2015). All branches were supported by ≥ 80% bootstrap values. The tree was rendered using the R package *ggtree* (Version 2.2.4) (Yu *et al*., 2017). Branches were colored according to the four phylogenetic groups (A through D) identified based on genome-wide single-nucleotide polymorphisms (SNPs). SNP groups A, B, C split into multiple clades, indicating rapid *vls* sequence divergence between closely related strains (Graves *et al*., 2013). Sequences within the multi-strain clades (e.g., within D group) did not separate into strain-specific subclades, suggesting frequent gene duplications and losses.

The *vls* sequences from the two SNP group D strains (JD1 and 156a) formed a monophyletic group consistent with the strain phylogeny. However, within this major clade the *vls* sequences did not separate into two strain-specific clades, a phylogenetic inconsistency that could be caused by gene duplications in a common ancestor followed by losses of duplicated copies or by incomplete lineage sorting, but unlikely by horizontal gene exchanges which would have introduced *vls* sequences from other SNP groups.

In contrast, the *vls* sequences from the strains belonging to the SNP groups A, B, and C all formed paraphyletic groups, each of which contained multiple clades highly divergent from one another than one would expect from the strain phylogeny (Fig 2). In the SNP group A, the *vls* sequences from the strain BOL26 formed a clade highly divergent from the *vls* sequences from the strains B31, PAbe, 64B, and ZS7. In the SNP group B, the *vls* sequences from three strains (WI91-23, N40, and 29805) formed three strain-specific clades. The *vls* sequences from strains belonging to the SNP group C were split into two clades, one consisting of the sequences from the strain 94a and the other consisting of sequences from the strains 72a and 118a.

As in the group D, the *vls* sequences within the SNP groups A, B, and C did not sort into strain-species clades, indicating frequent gene duplications, rapid gene losses, and fast sequence divergence within each of these phylogenetic groups. Indeed, it has been shown that the rapid sequence evolution of the *vls* cassettes was driven by adaptive differentiation evidenced by the accelerated nonsynonymous nucleotide substitutions (i.e., a high *dN/dS* ratio) (Graves *et al*., 2013).

### Evolutionary rates and molecular structure of *vls* cassettes

Rates of amino-acid substitutions are not uniform along the translated *vls* sequence, which consists of mostly fast-evolving variant regions (VRs) interspersed with six short conserved invariant regions (IR1-6) (Zhang *et al*., 1997). Here we quantified *vls* variability at individual amino-acid sites among the 13 *B. burgdorferi* strains using the 194 *vls* sequences including both the expressed and unexpressed cassettes. Conserved regions were detected by computing the relative evolutionary rate of each amino-acid site in the multiple sequence alignment, with the average variability score scaled to zero (Fig 3). Most residues in the IRs showed negative variability scores, indicating below-average evolutionary rates. The mean variability score for each IR was shown in Table 1. Among the IRs, IR1 was the least conserved followed by IR4. The IR2, IR3, and IR5 were conserved but relatively short. The IR6 was highly conserved at all 25 residues.

**Fig 3.**
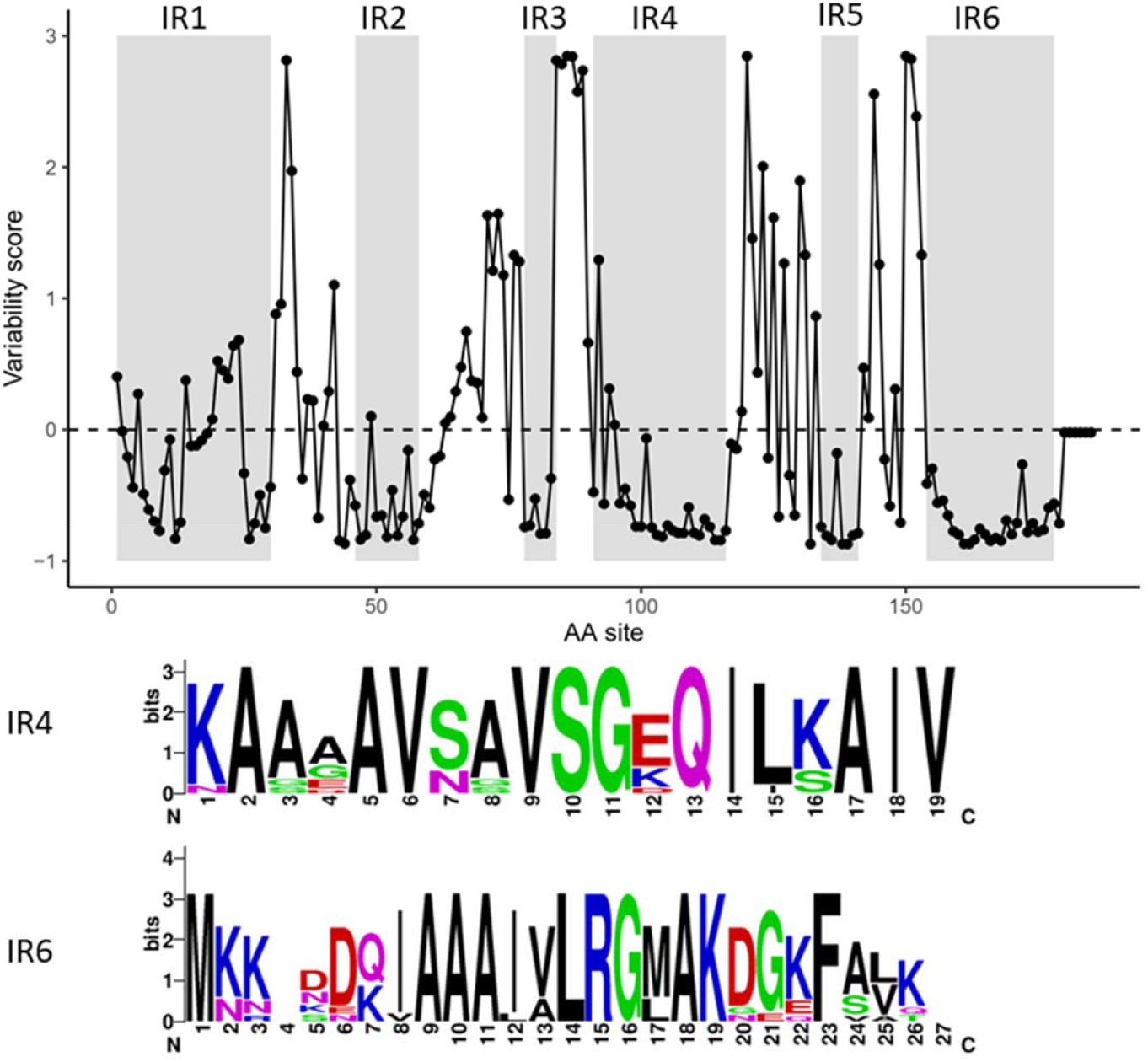
Site-specific evolutionary rates of *vls* cassettes (*Top*) Evolutionary rate, denoted as variability score (in the unit of standard deviation, *y-*axis), at each amino acid site was estimated by Rate4Site (Version 3.0.0) (Pupko *et al*., 2002) based on an alignment of translated sequences of 194 *vls* cassettes and the maximum-likelihood tree (Fig 2). The dashed line at 0 indicates the average evolutionary rate. The six IRs, showing generally lower-than-average rates, were shaded in gray. VlsE of B31-5A3 clone (GenBank accession U76405) was used as the reference for computation and annotation. (*Bottom*) SeqLogo images of IR4 and IR6 sequences, constructed based on one representative *vls* allele (translated) from each of the 12 *B. burgdorferi* genomes (Schutzer *et al*., 2011). Amino acid residues were colored according to physiochemistry. Letter heights correspond to information content in bits, a measure of site conservation (Crooks *et al*., 2004)

We further mapped the IRs to a published three-dimensional structure of the VlsE protein (from the strain B31) (Eicken *et al*., 2002) (Fig 4). All the IRs formed alpha helixes, as the ribbon model showed (Fig 4A). The space-filled model showed that the IR1, IR2, and IR4 were partially surface exposed while the IR2, IR5, and IR6 exhibited limited surface exposure (Fig 4B). The VlsE molecules likely form dimers on the spirochete cell surface (Eicken *et al*., 2002), which would further shield the invariant regions located on the monomer-monomer interface (Fig 4C and 4D). Nevertheless, the IR4 and IR6 are partially exposed at the membrane distal surface even in a dimerized form (Fig 4D).

**Fig 4.**
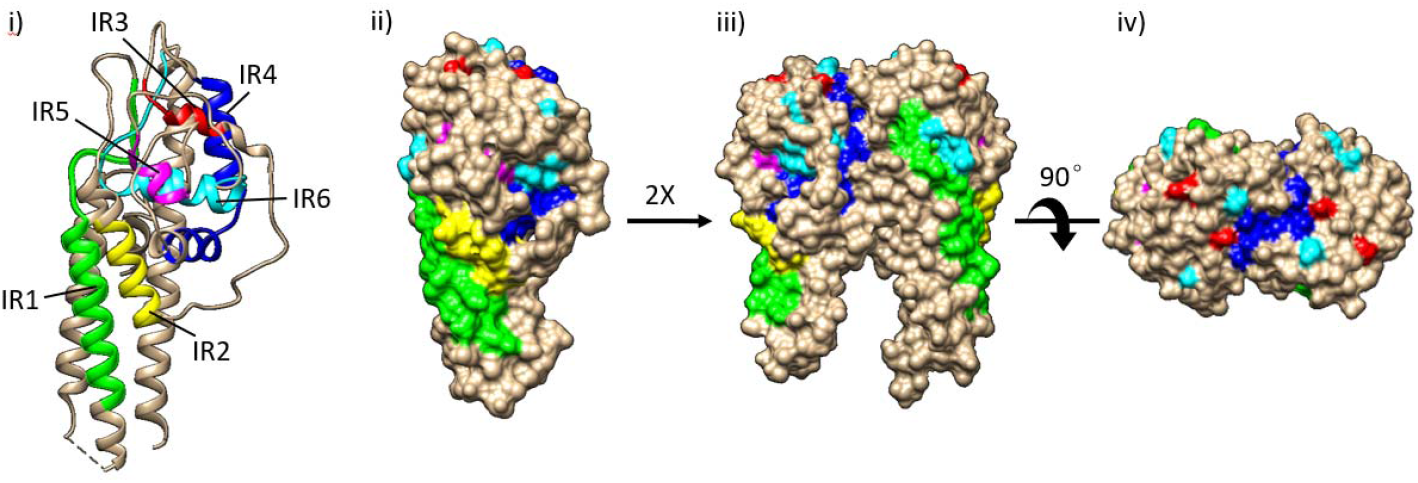
IR-highlighted three-dimensional structures of VlsE. Structure diagrams of VlsE protein from B31 were prepared in Chimera (Version 1.15) (Pettersen *et al*., 2004) based on the PDB file (accession 1L8W) (Eicken *et al*., 2002). IRs were highlighted in different colors (IR1 in green, IR2 in yellow, IR3 in red, IR4 in dark blue, IR5 in magenta, and IR6 in cyan). (*i*) Ribbon diagram showing that IRs tend to form alpha helices. (*ii*) Surface-filled diagram showing membrane surface exposure of IRs in monomeric form. (*iii*) and (*iv*) Dimerized structure models. The structures were oriented to show the membrane proximal part at the bottom (*i*, *ii*, and *iii*).

### Antigenicity of IRs against host sera

We measured antigenicity of the IRs with sera from human patients, white-footed mice, and immunized rabbits using ELISA. For the 46 human sera, the ELISA result showed an overall significant difference in the mean OD450 values among the antigens (*p* < 2.2e-16 by ANOVA) (Fig 5, left panel). Reactivities of the IR4 and IR6 peptides with the human sera were significantly higher than that of BSA (*p* = 2.87e-13 and 3.06e-13 by ANOVA, respectively), while reactivities of IR1, IR2, and IR5 were less significant (*p* = 0.034, 0.034, and 0.0019 by ANOVA, respectively) (Fig 5, left panel). Reactivity of VlsE was the strongest (*p* < 2.2e-16 by ANOVA). In addition, reactivities of the IR4 and IR6 with the human sera were weakly although significantly correlated with those of the VlsE (*p* = 7.6e-4 and *R^2^*=0.212 for IR4, *p* = 3.6e-3 and *R^2^*=0.158 for IR6, both by linear regression). There was no significant difference in reactivity between the early and late-stage patient samples (*p* = 0.8654 by *t*-test).

**Fig 5.**
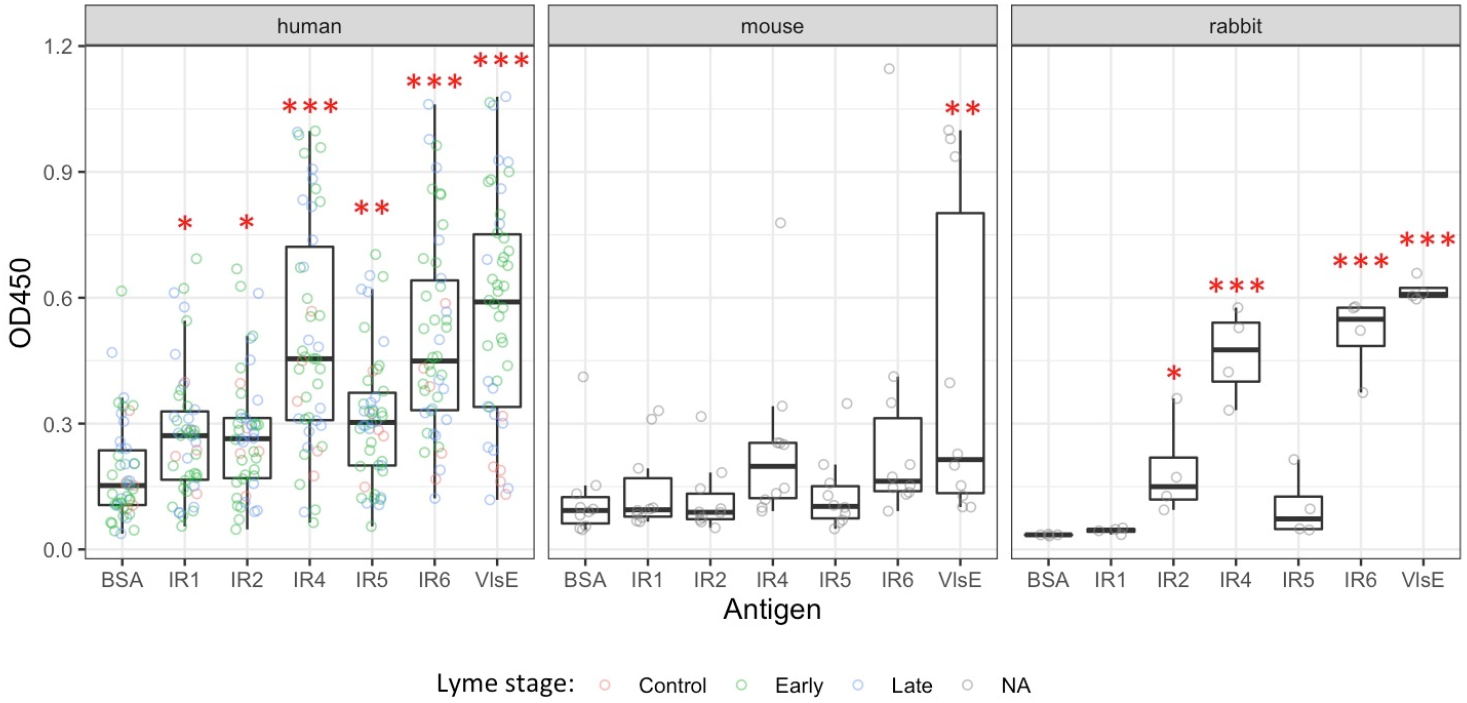
Antigenicity of conserved epitopes against host sera Antigenicity of five IRs (*x*-axis) was quantified with ELISA (see Material and Methods). Each IR peptide was tested for reactivity (OD450, *y*-axis) with host sera (represented by dots) from 46 human patients (*left*), 10 natural hosts (white-footed mouse, *Peromyscus leucopus*) (*middle*), and 4 New Zealand rabbits (*Oryctolagus cuniculus*) (*right*). Bovine serum albumin (BSA) was used as the negative control and the purified recombinant VlsE protein (of strain B31) as the positive control. Asterisks indicate significant differences between antigens and BSA by ANOVA analysis at varying degrees of confidence: “*” for 0.01< *p* < 0.05, “**” for 0.001 < *p* < 0.01, and “***” for *p* < 0.001.

Ten serum samples from whited-footed mice, the natural reservoir host of *B. burgdorferi* were tested. Reactivities of the IR peptides against the mouse sera showed little differences among the antigens (*p* = 0.0159 by ANOVA), with only VlsE showing a significant difference from the BSA control (*p* = 2.9e-3 by ANOVA) (Fig 5, middle panel). These results are consistent with findings of an earlier study which showed low antigenicity of the IR6 peptide in natural hosts relative to its antigenicity in humans (Liang *et al*., 1999).

Reactivities of the IRs against the four sera from four immunized rabbits showed a similar pattern as those against the naturally infected human (Fig 5, right panel). For example, VlsE, IR4, and IR6 peptides displayed the highest antigenicity (*p* = 6.8e-10, 1.2e-7, and 2.1e-8, respectively with an overall *p* = 2.2e-10 by ANOVA). Antigenicity of the IR1, IR2, and IR5 peptides against the rabbit polyclonal antibodies did not differ or differed weakly from that of BSA, the negative control (*p* = 0.855, 0.011, and 0.236 by ANOVA, respectively).

In sum, these ELISA results suggested that (1) anti-VlsE antibodies were present in patients throughout different stages of Lyme disease, (2) antibodies against the VlsE IRs were strongly present in naturally infected or artificially immunized non-reservoir hosts but minimally present in reservoir hosts, and (3) the IR4 and IR6 peptides were highly immunogenic conserved epitopes on the VlsE molecule in non-reservoir hosts relative to the IR1, IR2, and IR5 peptides. These results are consistent with conclusions of earlier studies on the antigenicity of VlsE and conserved epitopes, which established the use VlsE and the C6 peptide (derived from IR6) in both the standard and modified diagnostics tests of Lyme disease (Liang *et al*., 1999; Liang and Philipp, 1999, 2000; McDowell *et al*., 2002; Price, Dehal and Arkin, 2010; Branda *et al*., 2017; Pegalajar-Jurado *et al*., 2018; Lone and Bankhead, 2020).

Here we established that the IR4 peptide was as antigenic as the IR6 peptide. Indeed, both IR peptides reacted at a level similar to the reactivity of the whole VlsE protein with the sera from naturally infected and immunized hosts (Fig 5). The use of the highly conserved IR4 and IR6 as targets for theragnostic agents has the advantage that they are expected to exhibit antigenicity against a broad set of *B. burgdorferi* strains, with the potential to mitigate the challenge of strain-specific antigenicity of the highly variable antigens including VlsE and OspC (Bockenstedt *et al*., 1997; Bhatia *et al*., 2018).

### Identification and characterization of recombinant IR-specific monoclonal antibodies

Recombinant VlsE of the strain B31 was over-expressed, purified, and used to immunize New Zealand rabbit (Fig 6, gel image). IR-specific antibodies were identified via B cell sorting and by testing the reactivity of the supernatant of the 20 B cell lines against the five IR peptides with ELISA. We found that one cell line (1D11) bound specifically to the IR6 peptide and five cell lines (7C9, 15E2, 17A8, 28D3, and 42G10) specifically to the IR4 peptide in addition to their binding to the purified VlsE protein (Fig 6, bar plots). Supernatants of the remaining fourteen B cell lines reacted with the purified VlsE protein but not with the IR peptides, suggesting that the majority of B cell lines in the immunized rabbit expressed antibodies recognizing epitopes located on the variable and not the conserved regions.

**Fig 6.**
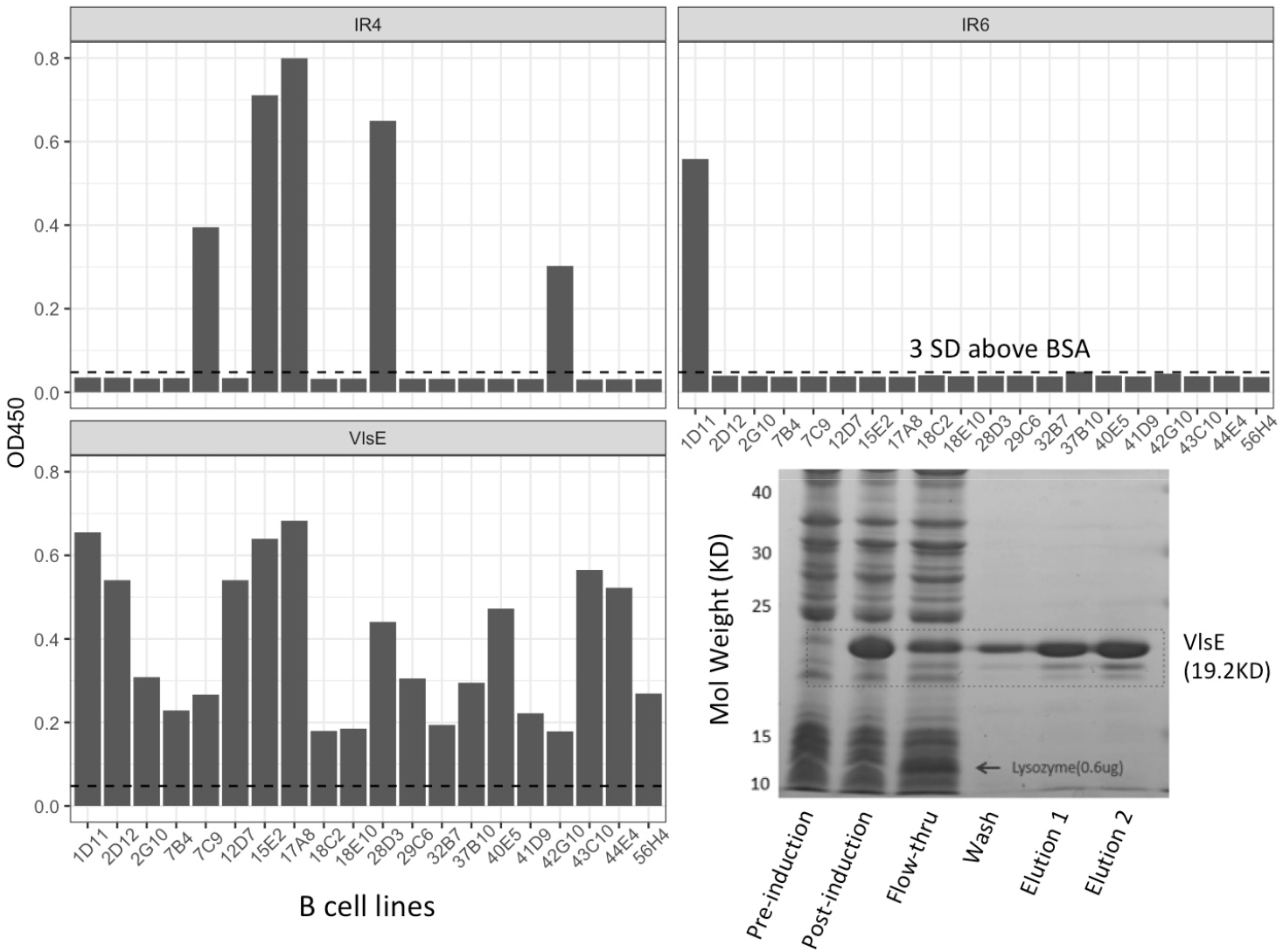
Identification of B cells producing IR-specific antibodies (*Bar plots*) Twenty VlsE-positive cell lines (*x*-axis) were selected with B cell sorting technique and tested against purified antigens with ELISA. Bovine serum albumin (BSA) was used as the negative control and the purified recombinant VlsE protein (of strain B31) as the positive control. An OD450 value (*y*-axis) greater than 3 standard deviations above the mean BSA reactivity (dashed lines) was considered to show significant antibody-antigen reactivity. Five cell lines expressing anti-IR4 antibodies and one cell line expressing anti-IR6 antibodies were identified. (*Image*) SDS-PAGE image of induction and purification of the recombinant B31 VlsE. Elution 1 and Elution 2 were combined into a single preparation with an estimated purity of ~65% and a concentration of ~5.0 mg/ml, which was subsequently used to immunize New Zealand rabbits.

One pair of the most abundant heavy chain and light chain variable region (V_H_ and V_L_) sequences in each of four IR-specific cell lines – including the anti-IR6 1D11 cell line and three top anti-IR4 cell lines – were identified by pyrosequencing and subsequently cloned and over-expressed. Specificity of the purified recombinant monoclonal antibodies (rMAb) were validated using ELISA. The initial rMAb cloned from the 1D11 cell line based on the most abundant V_H_ and V_L_ sequences was not reactive to the IR6 peptide as the supernatant of the cell line did. A new rMAb – based on the second most abundant V_H_ and V_L_ sequences – was re-cloned and over-expressed and reacted with the IR6 peptide strongly and specifically. The V_H_ and V_L_ sequences of the four IR-specific rMAbs and their binding characteristics were obtained by titration experiments (Fig 7).

**Fig 7.**
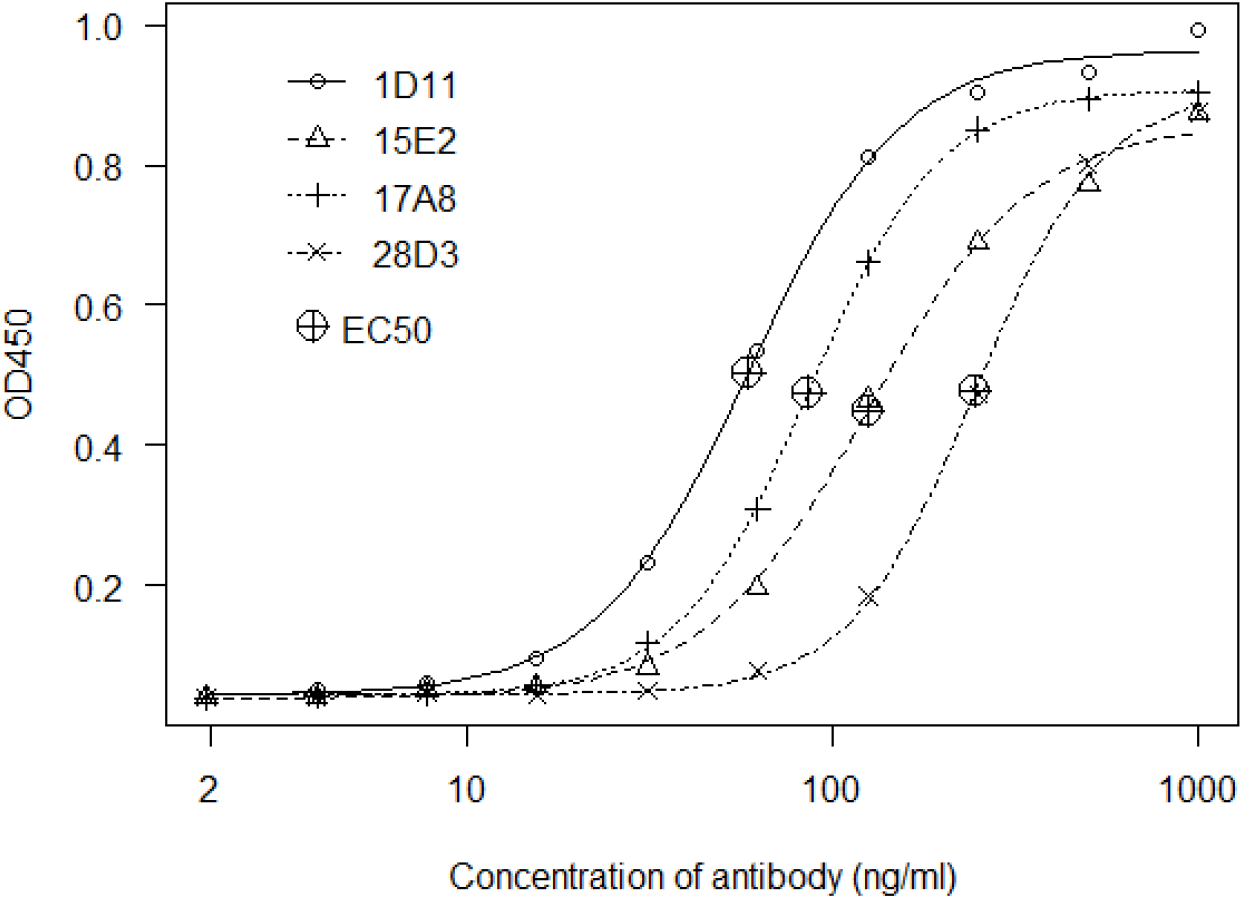
Binding characteristics of IR-specific monoclonal antibodies Serially diluted preparations of the four affinity-purified rMAbs were tested with ELISA against their respective IRs (1D11 against IR6; 15E2, 17A8 and 28D3 against IR4). The R package *drc* (Version 3.0-1) (Ritz *et al*., 2015) was used to estimate the effective concentration and to plot the titration curves. EC50 (effective concentration at 50% of the maximum activity) values were estimated from the fitted curves, with a lower EC50 indicating stronger antigen affinity.

## Discussion

### Rapid adaptive diversification of *vls* cassettes

The *vls* gene system in *Borreliella* was discovered based on gene sequence homology with the *vsp/vlp* (variable small and large proteins) system in *Borrelia* spirochetes causing relapsing-fever (Zhang *et al*., 1997; Norris, 2006). Since then, the molecular mechanism of segmental recombination between the expression site and the archival cassettes has been well characterized in *B. burgdorferi* B31, the type strain (Coutte *et al*., 2009; Verhey, Castellanos and Chaconas, 2019; Chaconas, Castellanos and Verhey, 2020). In parallel, genome-based comparative analysis of the *vls* system among *Borreliella* species and among strains of the same species uncovered rapid evolution in sequence, copy number, and genomic location of the *vls* cassettes (Glöckner *et al*., 2004; Graves *et al*., 2013; Schwartz *et al*., 2021).

In the present study, we showed pervasive phylogenetic inconsistencies between the *vls* gene tree and the genome-based strain tree, suggesting frequent gene duplications, gene losses, and gene exchanges, in addition to adaptive sequence evolution at the locus (Fig 2). The highly divergent *vls* cassette sequences between phylogenetic sister strains are reminiscent of the rapid amino-acid sequence diversification at the locus encoding the outer surface protein C (*ospC*), another immunodominant antigen of *B. burgdorferi* (Barbour and Travinsky, 2010). Protein sequences of major *ospC* alleles diverge in a strain-specific fashion with an average sequence identify of ~75.9% among *B. burgdorferi* strains in the Northeast US, due to a history of recombination among coexisting strains and diversifying selection driven by host immunity and possibly host-species preferences (Wang *et al*., 1999; Brisson and Dykhuizen, 2004; Haven *et al*., 2011; Di *et al*., 2021). In contrast, the coding sequences of the *vls* cassettes vary at a significantly higher level between the eight major sequence clusters (~56.3% average sequence identity), while varying at a high level between copies of the genome as well (e.g., 81.0% for B31, 76.5% for N40, and 76%.4 for JD1) (Fig 2). The much greater sequence diversity among the *vls* alleles than diversity among the *ospC* alleles indicates more rapid evolution driven by more intense immune selection at the *vls* locus.

As more *Borreliella* genomes are sequenced, the bioinformatics workflow including the customized web-based tool (http://borreliabase.org/vls-finder) established in the present study will facilitate large-scale automated identification of *vls* sequences and a quantification of the rates of gene duplication, losses, exchanges, and sequence divergence in this key adaptive molecular system in *Borreliella*.

### Immunogenicity of the IRs in non-reservoir hosts

The VlsE and its derivative C6 peptide (based on IR6) are key diagnostic antigens in serological tests of Lyme disease (Bacon *et al*., 2003; Marques, 2015; Branda *et al*., 2017). In the present study, we confirmed the predominant immunogenicity of IR6 in serum samples from human patients and VlsE-immunized rabbits (Fig 5). In contrast to an earlier study but consistent with another one (Liang *et al*., 1999; Chandra *et al*., 2011), the IR4 peptide showed as a similar level of antigenicity as the IR6 peptide in all three host species. Indeed, epitopes on IR4 might be more immunodominant than the IR6 epitopes in rabbits, as we obtained five anti-IR4 cell lines and only one anti-IR6 cell line out of a total of twenty randomly selected VlsE-reactive B cell lines (Fig 6). The IR1, IR2 and IR5 appeared to be barely immunogenic in reservoir as well as non-reservoir hosts (Fig 6).

Epitope mapping studies suggested that the IR6 may function as a single conformational epitope (Liang and Philipp, 2000). On an intact VlsE molecule (or its dimerized structure), the IR6 is almost entirely buried underneath the membrane surface and immunofluorescence assays demonstrated that the IR6 was inaccessible to antibodies on intact spirochetes (Liang, Nowling and Philipp, 2000; Embers *et al*., 2007; Elzbieta *et al*., 2016). It appears paradoxical that IR4 and IR6, two highly conserved and mostly buried regions on VlsE, contain immunodominant epitopes in human patients. Evolutionary arms races drive co-diversification of the antigen sequences in microbial pathogens along with the sequences of antigen-recognition proteins in vertebrate hosts through population mechanisms like negative frequency-dependent selection (Schierup, Mikkelsen and Hein, 2001; Haven *et al*., 2011; Papkou *et al*., 2019). Regions on antigen molecules shielded from host immune systems, like the IRs on VlsE, are not under such diversifying selection and thereby expected to be conserved in molecule sequences. The paradox resolves itself however when one considers that the IRs were indeed weakly immunogenic in the infected mice that belong to the natural reservoir species of *B. burgdorferi* (Fig 6). Indeed, as the whole VlsE molecule elicits significant antibody responses in the infected *P. leucopus* mice, such immunogenicity is likely due to epitopes on the variable regions as expected from the pathogen-host co-evolutionary arms race (McDowell *et al*., 2002) (Fig 6).

It is likely that *B. burgdorferi* is well adapted to infecting the natural hosts and able to maintain a high level of cell integrity including intact VlsE molecules on the cell surfaces throughout the infection cycle with a reservoir host. Indeed, *B. burgdorferi* expresses cell surface proteins binding specifically to proteins of the host complement system to down-regulate innate and adaptive host immunity (Kraiczy *et al*., 2006; Samuels, 2011; Hallström *et al*., 2013; Hammerschmidt *et al*., 2014). On an intact spirochete cell surface, the VlsE molecules can further shield other surface antigens from being recognized by antibodies (Lone and Bankhead, 2020).

In non-natural hosts such as humans and rabbits to which *B. burgdorferi* is poorly adapted, however, the pathogen may lose or diminish its ability to inhibit host immune responses and is thus more easily recognized by the host immune system. Upon cell disintegration and degradative processing of the surface antigens including VlsE by the major histocompatibility complex (HMC), the IR6 would be exposed along with other epitopes and elicit strong antibody responses. Since the IRs are conserved among the *vls* alleles and, unlike the VRs, their total amount remains stable during antigenic shift during infection, the IRs would result in stronger and more long-lasting host responses and become immunodominant in non-natural hosts including humans.

### Future work: needs for *in vitro* and *in vivo* validation

*B. burgdorferi* infection is characterized by a low number of colonizing spirochetes. It is difficult to directly detect the pathogen through culture or PCR approaches due to the extreme scarcity of the organism in infected hosts (Marques, 2015). Current diagnostic assays of Lyme disease, targeting the anti-VlsE or anti-C6 antibodies, do not distinguish between active and past infections (Wormser *et al*., 2013; Pegalajar-Jurado *et al*., 2018). Using recombinant monoclonal antibodies to directly detect the presence of spirochetes is a solution to the problem of low spirochete counts in human patients.

While we were able to obtain four IR4- and IR6-specific rMAbs, we recognize that the current study – which demonstrated their binding specificities to immobilized synthetic IR peptides using ELISA – is a key but only the first step. To validate the utility of these rMAbs as diagnostic and theragnostic agents, it is necessary to perform *in vitro* testing using cultured *B. burgdorferi* cells followed by *in vivo* testing using a mouse model of Lyme disease. We anticipate a number of biological and technical challenges during *in vitro* and *in vivo* validation testing of the rMAbs. For *in vitro* testing using cultured spirochetes, first, it is unclear if the rMAbs would bind VlsE anchored on the surface of live *B. burgdorferi* cells because of limited surface accessibility of the IRs at native conformations, even though the rMAbs reacted strongly with VlsE molecules fixed on a ELISA plate (Fig 2) (Liang and Philipp, 2000; McDowell *et al*., 2002). Molecular conformation is expected to differ among the synthetic peptides, the IRs in human sera, and the IRs on intact VlsE molecules anchored to the outer membrane of spirochete cells. Second, *B. burgdorferi* does not constitutively express a large quantity of VlsE during *in vitro* culture and supplementing the standard media with human tissue cells may be necessary to increase VlsE expression for *in vitro* validation of rMAb binding (Hudson *et al*., 2001). Third, both the IR4 and IR6 sequences vary slightly among *B. burgdorferi* strains despite high sequence conservation (Fig 3). The affinity of these rMAbs, which were raised using a single allelic variant (the B31 VlsE), is expected to vary among the *B. burgdorferi* strains. Effects of sequence variability to rMAb affinity could be quantified with ELISA using synthetic peptides representing the IR variants. Ideally, amino acid residues essential for the rMAb binding could be accurately pin-pointed with systematic epitope mapping (Chandra *et al*., 2011).

For *in vivo* testing of the rMAbs binding to the spirochetes, it is necessary to first to develop a live-imaging technology based on labeled rMAbs. Second, it is necessary to establish a murine model of Lyme disease in which laboratory mice (*Mus musculus*) are inoculated by needle or by infected ticks (Ivanova *et al*., 2009; Arumugam *et al*., 2019). For example, we plan to label the IR-species rMAbs with a radioactive isotope such as zirconium-89 and perform a positron emission tomography (PET) for the sensitive detection of trace quantities of spirochetes in experimentally infected mice.

## Author contributions

Li Li performed evolutionary analysis, protein purification, and ELISA. Li Li composed the initial draft. Lia Di developed the bioinformatics pipeline and web tool and participated in protein purification and ELISA. Saymon Akther participated in protein purification and ELISA and edited the manuscript. Brian Zeglis and Weigang Qiu conceived of, obtained funding for, and supervised the project. Weigang Qiu and Brian Zeglis revised the manuscript.

## Acknowledgments

This work was supported by the Public Health Service awards AI139782 (WGQ) from the National Institute of Allergy and Infectious Diseases (NIAID) and EB030275 (BZ and WGQ) from the National Institute of Bio-medical Imaging and Bioengineering (NIBIB) of the US National Institutes of Health (NIH). The content of this manuscript is solely the responsibility of the authors and does not necessarily represent the official views of NIH. Li Li and Saymon Akther are supported in part by the Doctoral Program in Biology of the Graduate Center, the City University of New York. We thank Dr. Christophe Sexton and Dr. Jeannine Petersen of the Center for Disease Control and Prevention (CDC) and Dr. Maria Gomes-Solecki (University of Tennessee Health Science Center) for providing sera samples from human patients and reservoir mice, respectively.

